# Disruption of endothelial stability directly impacts vascular neighboring cells in Hereditary Hemorrhagic Telangiectasia

**DOI:** 10.64898/2026.05.05.723108

**Authors:** Anna Sbalchiero, Luca Lambroia, Lorenzo Cassinelli, Roberta Carriero, Claudio Casali, Margherita Cavallo, Fabio Grizzi, Fabio Pasqualini, Mohamed AAA Hegazi, Sveva Introini, Fabio Sirchia, Carla Olivieri, Fabio Pagella, Leonardo Elia, Montserrat Climent

## Abstract

**BACKGROUND:** Hereditary hemorrhagic telangiectasia (HHT) is a genetic disorder caused by pathogenic variants in the endothelial TGFβ/BMP pathway, crucial for the vascular arterial-venous differentiation. Vascular defects result in fragile and malformed vessels. The precise mechanisms driving vascular network failure remain incompletely understood, complicating the design of targeted therapies.

**METHODS:** Nasal telangiectasias from HHT patients carrying variants in *ACVRL1* or *ENG* were used to perform scRNA-seq (2 *ACVRL1*- and 1 *ENG*-patient) and spatial transcriptomics (1 *ACVRL1* and 1 *ENG*) to uncover endothelial cells (EC) populations. Vascular characteristics within biopsies were evaluated using transmission electron microscopy (TEM) (1 *ACVRL1* and 1 *ENG*) and histological analyses (23 *ACVRL1* and 7 *ENG*), with particular attention to regions exhibiting varying degrees of damage.

**RESULTS:** Comparing our HHT tissues with healthy donor from the literature, we identified cellular heterogeneity within EC populations, revealing two distinct venous clusters: a stable, quiescent population (Mature Vein) and an activated, pro-inflammatory population (HHT Vein). The coexistence of these two clusters suggests cellular diversity within the biopsy, further validated by TEM and histology, revealing a juxtaposition of well-organized collagen and cellular architecture with severely disrupted, fibrotic regions. Moreover, cellular crosstalk analyses allowed us to identify critical ligands in ECs that interact with fibroblasts and mural cells. In particular, we found Midkine (MDK) lost in HHT Vein ECs with further validation *in vitro*, suggesting its potential role in cellular stability. Furthermore, spatial transcriptomics allowed to further uncover pathologic phenotypes in cells neighboring HHT Vein ECs.

**CONCLUSIONS:** HHT biopsies exhibit localized inflamed and fibrotic vascular areas with the presence of different transcriptional sub-populations of EC. Within the same tissue, stable and activated ECs can be distinguished. The pathologic-like EC cluster, present exclusively in the HHT samples, may contribute to vascular leakage through the loss of important ligands involved in cellular communication.

## Introduction

Hereditary hemorrhagic telangiectasia (HHT) is a rare genetic vascular disorder characterized by multiple abnormal blood vessel connections that bypass the capillary bed, leading to the direct arteriovenous shunts, causing the development of fragile, leaky vessels^1^. These alterations are known as telangiectasia, when small and located on the skin and mucocutaneous surfaces ^2^, and arteriovenous malformation (AVMs), when larger and present in internal organs, such as gastrointestinal tract, liver, lung, and brain ^3^. These malformations are classified by vessel type and flow dynamics: capillary, lymphatic, venous, and combined forms are slow-flow, whereas arteriovenous malformations (AVMs) are fast-flow lesions. Early vascular changes involve focal dilation of post-capillary venules, which progressively enlarge and develop a disproportionate smooth muscle cell (SMC) layer ^4^.

HHT is caused by heterozygous, loss-of-function pathogenic variants in components of the endothelial Transforming Growth Factor β (TGFβ) / Bone Morphogenic Protein (BMP) signaling pathway, which is crucial for vascular integrity and function ^5^. Pathogenic variants are frequently found in *ENG* and *ACVRL1*, which encode a co-receptor (ENG) and a type I receptor (ALK1) of the endothelial BMP pathway, responsible for HHT1 (MIM #187300) and HHT2 (MIM #600376), respectively^6,7^. The TGFβ/BMP signaling pathway is essential for endothelial homeostasis therefore, dysregulation of this axis profoundly impacts endothelial processes, including proliferation, migration, and vessel maturation^8–10^.

In endothelial cells (ECs), BMP9/10 bind the ENG-ALK1 receptor complex inducing phosphorylation of SMAD1/5/8 and activating transcriptional programs that promote endothelial quiescence, flow responsiveness, and proper arterial–venous identity^5,11,12^. Indeed, loss-of-function mutations in *ENG* or *ACVRL1* diminish BMP9/10 signaling, impair flow-mediated responses, and shift EC behavior toward excessive migration, abnormal angiogenesis, and heightened VEGF sensitivity^13,14^. Furthermore, reduced ENG–ALK1 activity drives signaling toward the TGFβ/ALK5–SMAD2/3 axis, which promotes endothelial inflammation, upregulation of adhesion molecules, and induction of endothelial-to-mesenchymal transition (EndoMT) ^15^. EndoMT contributes to extracellular matrix (ECM) remodeling and fibrosis, including collagen deposition, which are all histological features frequently observed in HHT vascular lesions^15^. Thus, the imbalance between protective BMP9/10–ALK1 signaling and TGFβ-mediated inflammatory and fibrotic responses provides a mechanistic link between HHT-causing mutations and vascular malformation development. An increased body of work has uncovered the disruption of signaling pathways, such as PI3K-AKT in ALK1-HHT biopsies causing excessive endothelial proliferation ^16,17^ and development of fibrosis and inflammation within HHT tissues^18^.

Despite this growing knowledge, the precise mechanisms driving vascular network failure remain incompletely understood, complicating the design of targeted therapies ^13,19^. Available therapies have several limitations, including poorly defined mechanisms of action, uncertain off-target effects, and treatment resistance or relapse in some patients. Therefore, identifying specific vascular defects, while accounting for the underlying cellular complexity, will be essential for guiding more precise and effective, individualized therapeutic strategies for patients with vascular malformations.

In this study, we characterize the cellular complexity of telangiectasias from nasal biopsies of HHT patients, with a particular focus on the transcriptional heterogeneity of EC populations. We identified a distinct EC cluster exhibiting pathological features of endothelial activation, inflammation, and EndMT, processes closely associated with fibrosis and collagen deposition. Together, these findings contribute to the validation and establishment of a framework for understanding how endothelial dysfunction shapes the pathogenesis of HHT and set the stage for identifying molecular vulnerabilities that could be targeted therapeutically.

## Methods

### Ethical agreement and Patient biopsies

Patients were recruited from the HHT Reference Unit at IRCCS Fondazione San Matteo in Pavia and were considered eligible for enrollment if they met at least three Curaçao criteria^6,20^. All patients signed the informed consent for nasal biopsy in accordance with the Clinical Research Ethics Committee requirements from IRCCS San Matteo (Ethical Committee protocol HHT-BersagliMolecolari_GJC23034). Patient data were anonymized, and each individual was assigned an alphanumeric code. Baseline epidemiological and clinical characteristics of the patients are reported in Table 1.

**Table 1.**
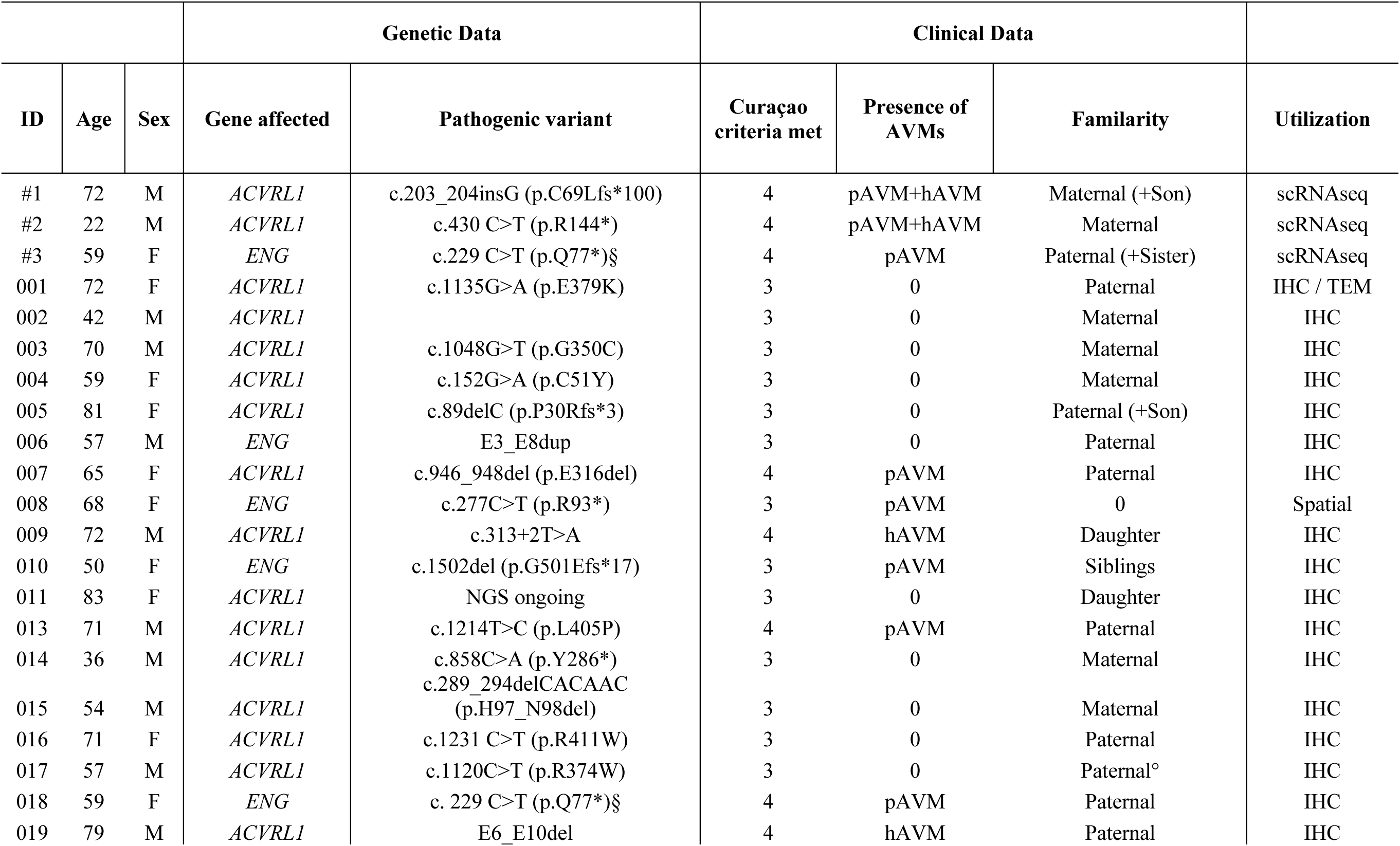
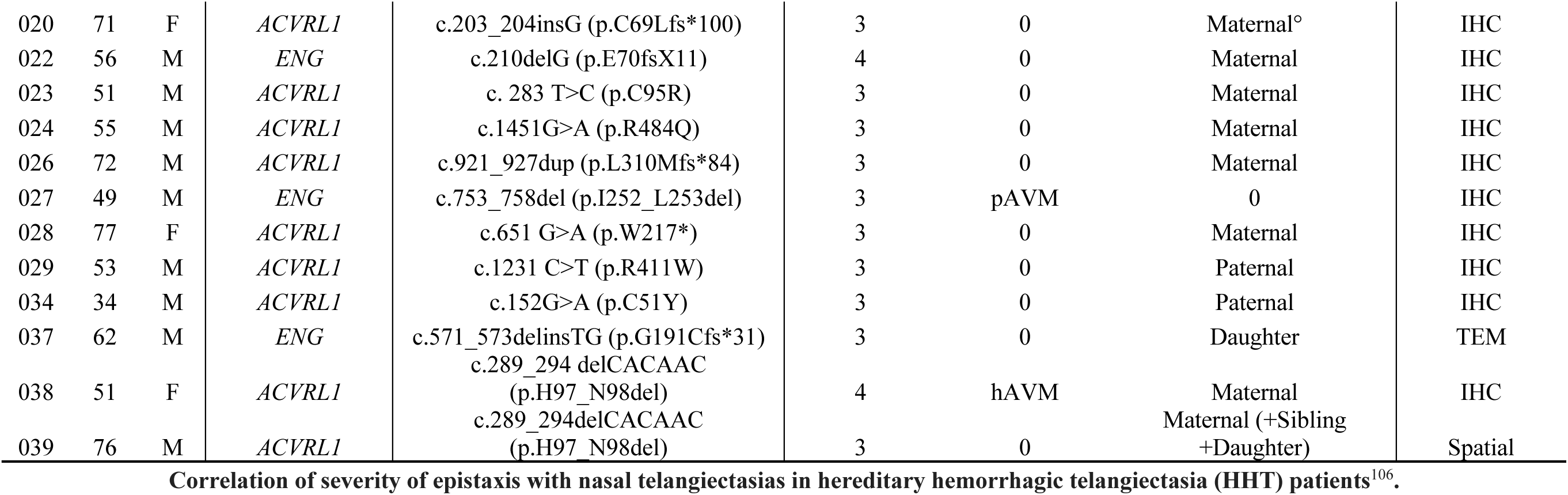
Clinical and genetic characterization of HHT patients.

### Immunofluorescence

FFPE tissues were sectioned at 3 µm. Sections were deparaffinized in xylene two times, ten minutes each and rehydrated through a graded ethanol series (100%, 100%, 90%, 70%, one minute each), following by rinsing in distilled water. Antigen retrieval was performed by immersing the slides in a retrieval solution (EDTA, 0.5 M, pH 8.0, Molecular Biology Grade, DEPC-Treated Merck, Germany, dilution 0.05% in distilled water) and heating in a microwave. Multiplex immunofluorescence assay was performed using the Opal Polaris™ 7-color manual IHC Kit (NEL861001KT) following the manufacturer’s instructions. Primary antibodies against FosB (Cat # MA5-15056, Invitrogen, USA, dilution 1:200), IL1R1 (Cat # PA5-119223, Invitrogen, USA, dilution 1:200) and PTGDS (Cat # 10754-2-AP, Proteintech, USA, dilution 1:250) were conjugated with Opal 520, Opal 620 and Opal 690 fluorophores, respectively. Nuclei were counterstain with DAPI (1:5000).

### Computer-assisted vessel image analysis

#### CD34/ACTA Immunofluorescence (IF)

For multiplex immunofluorescence analysis, all CD34- and ACTA-stained tissues were digitized using a Zeiss Axioscan.Z1 slide scanner (Zeiss, Germany) at 20× objective magnification. Multi-channel quantitative analysis was then performed with Angio Lab v1 (Histology Core, IRCCS Humanitas Research Hospital, Milan, Italy). In all samples, CD34 staining was used as the vascular reference channel for the identification and segmentation of CD34⁺ vascular structures. ACTA signal intensity was quantified within the CD34-defined vascular regions. For each segmented CD34⁺ vessel, the software automatically extracted ACTA signal metrics, including mean ACTA intensity and standard deviation (SD) within the CD34⁺ area. Vessels were subsequently classified based on marker expression as CD34⁺/ACTA⁻ or CD34⁺/ACTA⁺, and their relative proportions were calculated. Vessels were further stratified into small and large groups using the same area threshold of 700 µm².

#### Nearest Neighbor Index (NNI)

To assess whether vascular spatial organization deviated from complete spatial randomness, spatial pattern analysis was performed using the Nearest Neighbor Index (NNI). The analysis was applied to both CD34⁺/ACTA⁻ and CD34⁺/ACTA⁺ vessel populations, using the following formula: *NNI = observed mean distance / expected mean distance*

NNI values < 1 indicate spatial clustering, values ≈ 1 are consistent with a random spatial distribution, and values > 1 reflect spatial dispersion or regularity.

### Single cell RNA-sequencing library preparation

Three nasal biopsies were collected during Argon plasma coagulation surgery^21^ from patients carrying *ACVRL1* (n=2) and *ENG* (n=1) variants. Biopsies were minced and placed in digestion solution with dispase (Cat#SCM133, Merck) and collagenase IV (Cat# LS004189, Worthington Biochemical Corpotation) for 45 min at 37 degrees. After, digestion was stopped with pure FBS and cell suspension was centrifuged. Suspension was cleared from red blood cells by lysis with ACK lysis buffer (Cat# A1049201, Life Technologies) and washed with 1x PBS. Finally, cells were suspended in 1x PBS and counted using using trypan blue staining. Cell viability was > 95%. 10000 cells were loaded onto 10X Genomics Chromium Controller for droplet generation.

Reverse transcription and library preparation were performed with the Chromium Next GEM Single Cell 3ʹ GEM according to 10X Genomics’ standard protocol. Libraries sequenced in the NextSeq1k2k from Illumina.

### Single cell RNA-sequencing analysis

Single-cell RNA-seq data were processed using Cell Ranger v6.1.1 (10x Genomics), aligning sequencing reads to the human GRCh38 reference genome. Raw count matrices were analyzed in R v4.0.5 using DropletUtils v1.14.2 and scater v1.24.0, together with scran v1.34.0 and scDblFinder v1.20.2. Empty droplets were identified and removed using emptyDrops (FDR ≤ 0.05). Outliers for UMI counts, gene counts and mitochondrial or ribosomal read percentages were filtered based on median absolute deviation (isOutlier, nmads = 3) and cells with fewer than 500 detected genes were excluded. Genes expressed in fewer than 10 cells with at least one UMI were removed. Putative doublets were identified by computing doublet density and removed. Filtered count matrices were analyzed by Seurat v5.0.0 and cells were further filtered using conservative thresholds (nFeature_RNA ≤ 8000 and percent.mito < 20%). Public scRNA-seq biopsy samples from nasal, tracheal and intermediate airway tissues described by Deprez et al.^22^ (Dataset ID: https://ega-archive.org/studies/EGAS00001004082) were included as healthy controls, and cluster annotation was performed based on correspondence with healthy reference populations and expression of top marker genes identified using FindAllMarkers, together with established markers from the literature. Data integration across samples was performed using Harmony v1.2.0 via the Seurat-Harmony wrapper followed by principal component analysis, graph-based clustering and UMAP visualization. Endothelial clusters were isolated and re-clustered independently, with normalization, dimensionality reduction, and clustering steps repeated. Endothelial subtypes were then annotated based on established marker genes reported in the literature^23,24^. Differential gene expression analyses were performed using Seurat’s FindMarkers function with the MAST statistical framework, both across all annotated cell types and in focused comparisons between endothelial subtypes and healthy Mature Vein; for all comparisons against Mature Vein, only healthy Mature Vein cells derived from nasal biopsy samples were used as the reference population. P-values were corrected for multiple testing using the Benjamini–Hochberg method, and genes with adjusted p-values < 0.05 were considered differentially expressed. Gene set enrichment analyses of differential gene expression results were performed using clusterProfiler v4.14.6, interrogating the Gene Ontology Biological Process and HALLMARK database. Regulon inference in endothelial cells was performed using SCENIC v1.3.1, and regulon activity at the single-cell level was quantified using AUCell v1.28.0. Cell–cell communication between HHT Vein, Mature Vein and stromal cell populations (fibroblasts, smooth muscle cells and pericytes) was inferred using CellChat v2.1.2. The human CellChat ligand–receptor database was used, and the standard CellChat workflow was applied; to increase biological specificity, only ligand–receptor pairs for which at least one component was identified as differentially expressed in the corresponding FindMarkers analyses between patient’s derived HHT Vein and healthy nose biopsies Mature Vein. All visualizations were generated using ggplot2 v3.5.1, either directly or through Seurat and CellChat plotting functions.

### Tissue Processing and Spatial Transcriptomics Sequencing

For this analysis, one *ACVRL1* and one *ENG* biopsies were used. Spatial transcriptomics Gene Expression Kit (BMKMANU, ST03002) and Tissue Optimization Kit (BMKMANU, ST03003) were used according to the manufacturer instructions. Each capture area of the gene expression slide (6.8 x 6.8 mm^2^) contains 4,140,000 barcoded spots that are 2.5 μm in diameter (3.5 μm center to center between spots) providing an average of 7 to 37 spots under a cell. The frozen tissue was embedded in Optimal Cutting Temperature compound (OCT, Sakura Tissue-TEK) on dry ice and then stored at -80°C. The Tissue Optimization Kit was first performed according to the manufacturer’s instructions, and the fluorescent footprint was imaged using a Metafer Slide Scanning Platform (Pannoramic MIDI) to select the optimal permeabilization time. OCT blocks were cut with a pre-cooled cryostat at 10 μm thickness, and sections were transferred to fit the 6.8 x 6.8 mm2 oligo-barcoded capture areas on the BMKMANU S3000 Gene Expression Slide (BMKMANU, ST03009). The Gene Expression Slide with tissue was fixed and stained with hematoxylin and eosin (H&E) and imaged using the Pannoramic MIDI microscope at 40X magnification. [Nuclei staining was also performed and imaged using fluorescent and transflective setting for subsequent cell segmentation.] The Pannoramic MIDI was used to acquire tile scans of the entire array and merge images. Permeabilization was then carried out using the optimal time, followed by reverse transcription. Sequencing libraries were then prepared according to the manufacturer’s instructions using the Library Construction Kit (BMKMANU, ST03002-34). Libraries were sequenced on a NovaSeq X Plus platform (Illumina) at a minimum sequencing depth of 150,000 read pairs (PE150) per spatial spot.

#### BMKMANU S3000 Spatial Transcriptomics Data Analysis

Raw sequencing data were firstly processed using the BSTMatrix pipeline, which performs spatial barcode demultiplexing, alignment to the human reference genome (GRCh38, release 95), and unique molecular identifier (UMI)-based quantification to generate spot-level gene expression matrices. Automated image-based cell segmentation and tissue region identification were integrated to assign transcripts to spatial locations and reconstruct cell boundaries. Across the two analyzed samples, a total of 40,518 and 17,636 cells were identified, respectively, from 1,162,862 and 303,270 spatial spots. A cell segmentation strategy was applied, allowing transcripts to be assigned to individual cells and enabling spatial analysis at single-cell resolution. The resulting spatial gene expression matrices were exported and converted into R-compatible objects (RDS format) using the BSTMatrix tool, enabling downstream analysis with Seurat R’s package.

#### Spatial transcriptomics Clustering analysis

All downstream analyses were performed in R (v4.4.3) using Seurat (v5.0.3). After data normalization and variance stabilization using SCTransform on the Spatial assay, dimensionality reduction was performed by principal component analysis (PCA) on the SCT assay. The first 30 principal components were used to construct a shared nearest-neighbor graph (FindNeighbors, dims 1–30), and unsupervised clustering was performed with FindClusters. Multiple clustering resolutions (0.0–1.0, step 0.1) were explored, and cluster stability across resolutions was evaluated using the clustree package. Cluster annotation was performed by integrating canonical marker gene expression and signature-based scoring. Cell-type transcriptional signatures were grouped into higher-order macro-populations (epithelial, olfactory, lymphoid, myeloid, granulocyte/mast, stromal, vascular). For each macro-population, a group score was calculated per cluster. Spatial coherence of selected group scores, including vascular signatures, was further evaluated using spatial feature visualization. To further resolve heterogeneity within the vascular compartment, the cluster enriched for endothelial and vascular signatures was subset and reanalyzed using the same workflow.

#### Spatial transcriptomics neighbor analysis

To characterize the spatial microenvironment surrounding vascular structures, a spatial neighbor analysis was performed. Clusters of interest (annotated as HHT or mature vein) were defined as points of interest (POIs). A neighborhood radius of approximately 100 µm was selected and converted into pixel units using image-specific scaling factors to ensure consistency with the spatial coordinate system.

For each POI, neighboring cells within the defined radius were identified using a radius-based nearest-neighbor search implemented in the RANN package. Unique neighboring cells were aggregated across all POIs, excluding the POIs themselves, thereby defining the perivascular neighborhood.

The resulting cells were subsequently clustered, and cell identities were assigned based on marker gene expression. Differential gene expression between neighbor clusters was assessed using the *FindMarkers* function.

### Statistical Analyses

Data are expressed as mean ± SD (standard deviation), calculated by GraphPad Prism 10 software. Statistical analyses were performed on at least three independent experiments. Individual values were scanned for the presence of outliers using ROUT test. For the assumption of parametric tests, the Gaussian distribution of values through Shapiro-Wilk normality test or Kolmogorov-Smirnov test were evaluated. Parametric unpaired Student’s t test or Mann-Whitney test were applied in the case of two groups of analysis (two-tailed). In the case of more than two groups of analysis, univariate ANOVA (1-way ANOVA corrected for multiple comparisons test) in presence of one independent variable or the factorial design ANOVA (2-way ANOVA corrected for multiple comparisons test) to examine the effect of two factors on a dependent variable were used. Statistical significance was defined as P ≤ 0.05.

## RESULTS

### Cellular populations in human HHT nasal telangiectasias

To characterize the cellular landscape of human HHT lesions, we performed single-cell RNA-sequencing (scRNA-seq) on three nasal telangiectasia biopsies from three HHT patients (2 *ACVRL1*-patients and 1 *ENG*-patient, Table 1). As controls, we included previously published scRNA-seq data from healthy human airway biopsies obtained from the study by Deprez et al ^22^. We profiled a total of 22175 cells from HHT telangiectasias and 46791 cells from control samples. Unsupervised clustering revealed 18 distinct cell populations, each characterized by unique transcriptional profiles (Figure 1A). Cell identities were assigned based on canonical markers genes ^23^ (File S1), and the top three genes defining each cluster are shown in Figure S1A. As expected for nasal mucosa, epidermal-related cells, including basal, suprabasal, cycling basal, secretory, serous, and ciliated cells, represent a large fraction of the dataset. After evaluating the cellular proportions (Figure 1B), we observed that inflammation-related cells, such as B cells, Basal, Dendritic, Multiciliated, Smooth muscle and Suprabasal cells; showed a significant increase in frequency in HHT biopsies, as expected in damaged tissue ^16,25^. Interestingly, vascular cell populations were also notably increased in HHT lesions compared to heathy tissue. These included fibroblasts (FBs), enriched for *SFRP2* (Secreted Frizzled-Related Protein 2), *LRRC15* (Leucine-Rich Repeat Containing 15), *SFRP4* (Secreted Frizzled-Related Protein 4); pericytes (PCs), expressing *FAM162B* (Family With Sequence Similarity 162 Member B), *KCNJ8* (Potassium Inwardly Rectifying Channel Subfamily J Member 8), *HIGD1B* (HIG1 Hypoxia Inducible Domain Family Member 1B); and smooth muscle cells (SMCs), marked by *CNN1* (Calponin 1), *ACTG2* (gamma-2 actin) and *PLN* (phospholamban). Interestingly, ECs, identified by the expression of *ECSCR* (Endothelial Cell Surface Expressed Chemotaxis and Apoptosis Regulator), *ACKR1* (Atypical Chemokine Receptor 1, formerly DARC) and *CLDN5* (Claudin-5), were present at similar proportions in healthy and HHT samples. We evaluated the up- and down- regulated pathways in these vascular populations by comparing HHT patients to healthy donors and observed an overall activation of pathways related to TGFβ response, Wnt signaling, cell adhesion, response to oxygen levels, wound healing, as well as pathways involved in vascular development, including vasculogenesis and endothelium development (Figure 1C). Interestingly, ECs from HHT specimens showed activation of pathways related to cell number regulation, indicating a possible cell-autonomous mechanism maintaining the stable proportion of ECs between healthy and HHT samples. Other enriched pathways included response to hormones, oxidative stress, and inflammation. On the other hand, down-regulated pathways were mainly related to oxidative stress homeostasis and lipid metabolism.

**Figure 1.**
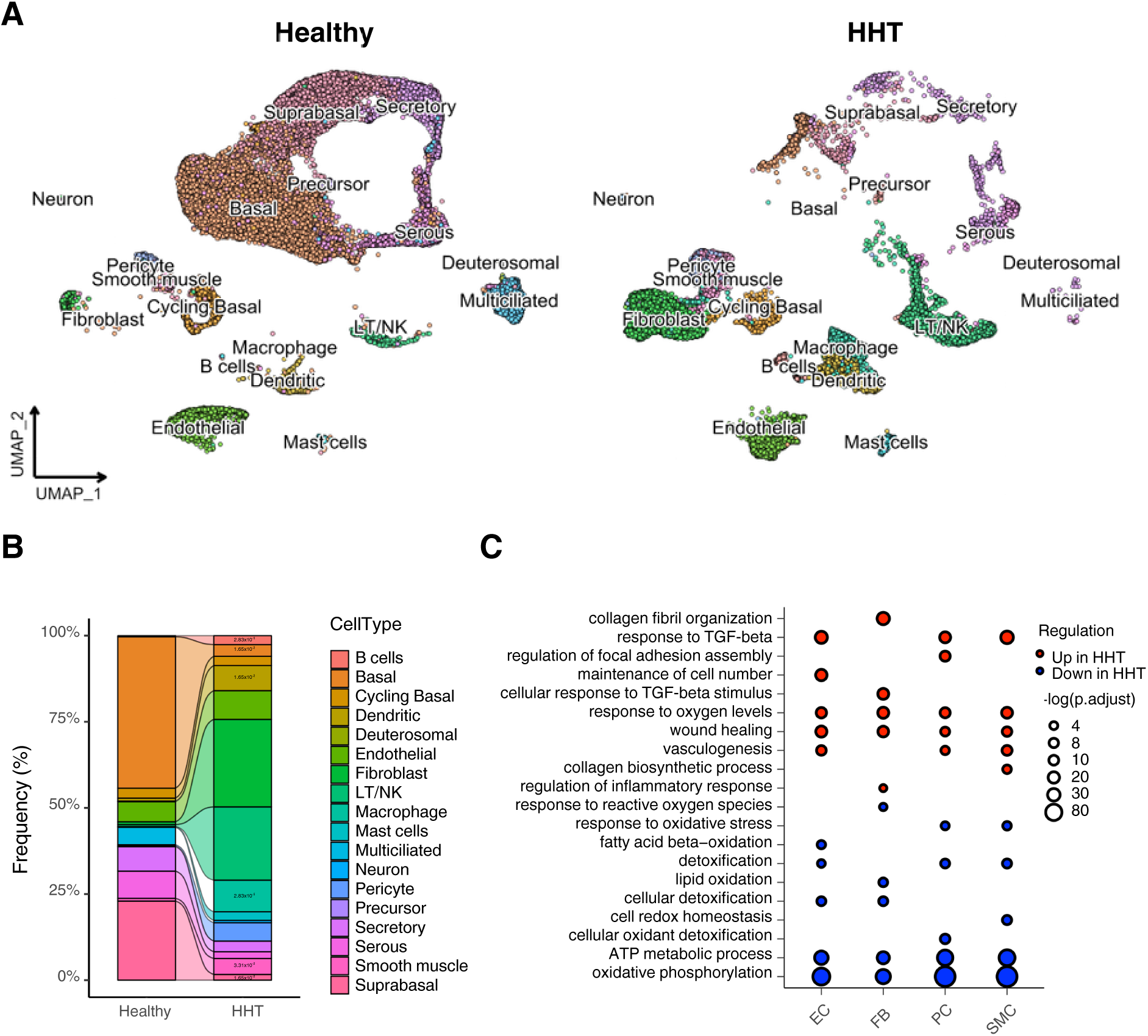
Identification of cellular populations in nasal HHT telangiectasias. **A**, UMAP plots of all cellular populations showing differences between Healthy donors (left) and HHT biopsies (right). **B**, Histograms displaying frequency of the different cellular populations, in percentages (%), in Healthy donors and HHT patients. Numbers indicate p-value statistically different cellular frequency between Healthy and HHT samples. **C**, Dot blot showing differential pathways between different populations: EC (endothelial cells), FB (fibroblasts), PC (pericytes) and SMC (smooth muscle cells). Up-regulated pathways (red) and down-regulated pathways (blue) in HHT compared to Healthy donors.

Overall, this initial assessment indicates that HHT lesion reshape the tissue cellular composition primarily by expanding inflammatory and vascular stromal compartments. Although ECs are well known to play a central role in the diseases, their stable abundance in the scRNA-seq data highlights the importance of dissecting their molecular heterogeneity and state-specific alterations.

### Endothelial heterogeneity in human HHT nasal telangiectasias

The roles of *ACVRL1* and *ENG* in EC biology have been widely investigated ^26–32^ and pathogenic variants in these genes are directly implicated in the AVMs and vascular fragility in HHT ^9^. Given the central role of ECs in HHT development and their active phenotype, we performed a detailed characterization of EC transcriptional states.

We first assessed the expression of the causative genes specifically within the EC population of each biopsy. As expected, *ACVRL1* and *ENG* expression was markedly reduced in biopsies carrying pathogenic variants in *ACVRL1* (#1 and #2) and *ENG* (#3), respectively (Figure 2A). Because these genes are integral components of the TGFβ/BMP signaling pathway, we next evaluated the expression of major pathway-associated genes. We observed a pronounced dysregulation of the TGFβ/BMP signaling in EC from HHT samples compared to ECs from healthy donors (Figure 2B), with genes active in healthy ECs being suppressed in the HHT ECs and vice-versa, consistent with the overall pathway imbalance during HHT pathology.

**Figure 2.**
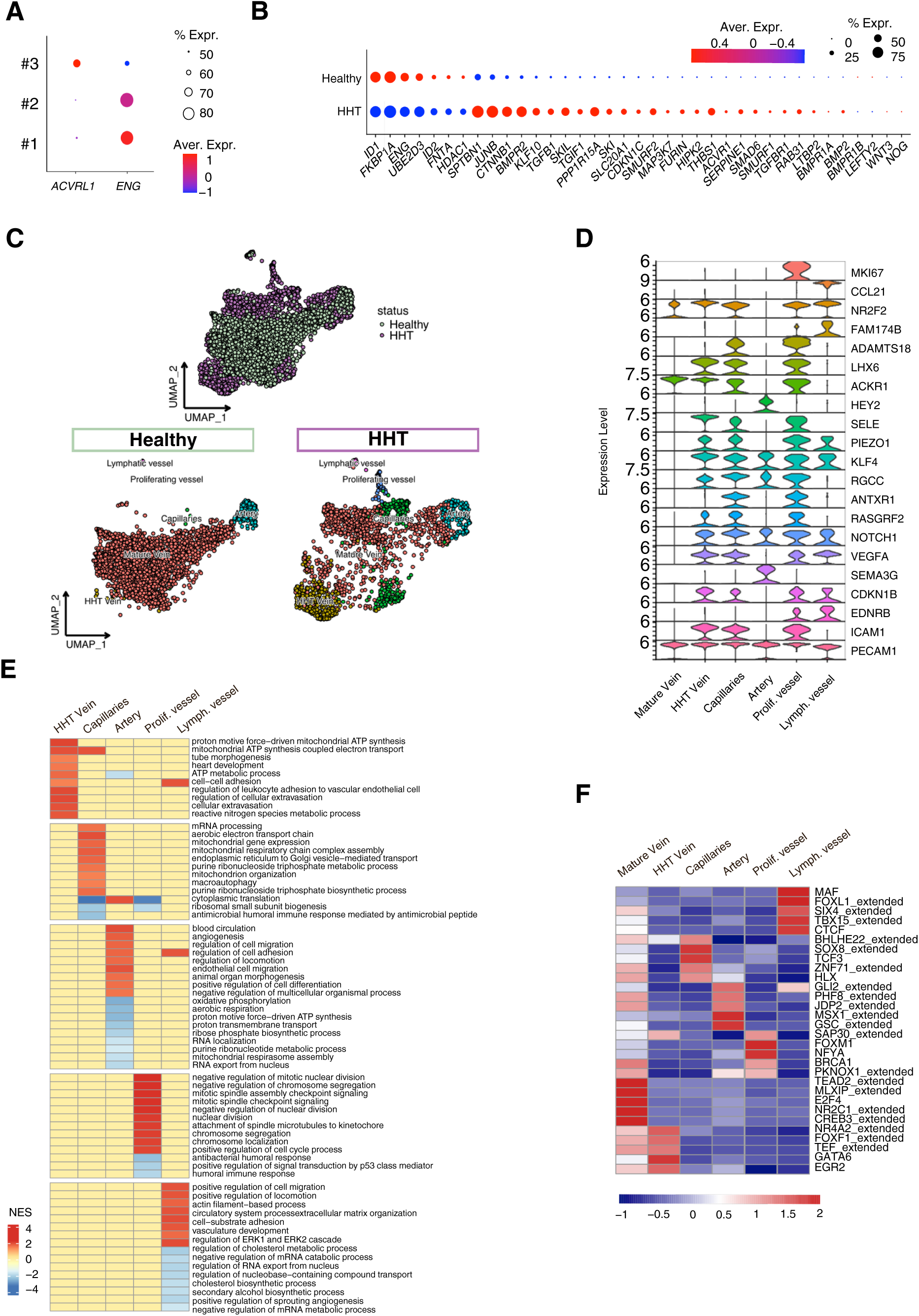
Identification of endothelial sub-populations in nasal HHT biopsies. **A**, Dot blot showing RNA expression levels of *ACVRL1* and *ENG* from each biopsy (#1 and #2, *ACVRL1*; #3 *ENG*) in all EC populations. Color scale: red, high expression; blue, low expression. **B**, Dot blot presenting the expression levels and frequency of the selected genes involved in the TGFβ/BMP pathway among Healthy and HHT patients. Color scale: red, high expression; blue, low expression. **C**, UMAP plot displaying the EC distribution Healthy and HHT merged (top) and distinguishing between Healthy (bottom left) and HHT (bottom right) samples. **D**, Violin plots of gene expression of EC lineage markers identifying the different sub-clusters of ECs. **E**, Heat map of the 5-top enriched pathways from each EC sub-cluster form HHT samples compared to the Healthy samples. All relative to Mature Vein cluster. Color scale: red, high expression; blue, low expression. **F**, Heat map with the top transcription factors active in each EC sub-population. Color scale: red, high expression; blue, low expression.

Because ECs from HHT biopsies showed an overall switch in phenotype towards pathogenic pathways (Figure 1C and 2B), we then proceeded to dissect EC heterogeneity within telangiectasias. The analysis of the overall EC population from healthy and HHT samples revealed substantial transcriptional differences and a marked redistribution of EC populations in HHT tissues (Figure 2C). To refine these observations, we performed EC sub-clustering annotating vessel subtypes according to gene expression of markers from literature ^23,24^. This analysis identified six EC sub-clusters, the UMAP shows the distribution of the different EC sub-clusters in healthy and HHT samples (Figure 2C), while the violin plot shows the main expressed markers for each sub-cluster (Figure 2D). In healthy samples, the principal EC clusters correspond to: a) Mature vein, expressing venous markers such as *ACKR1* and *NR2F2* (Nuclear Receptor Subfamily 2 Group F Member 2); b) Artery, defined by *HEY2* (hes related family bHLH transcription factor with YRPW motif 2), *NOTCH1* (Neurogenic locus notch homolog protein 1) and *SEMA3G* (Semaphorin 3G); c) Capillaries, characterized by *RGCC* (Regulator Of Cell Cycle), *ANTXR1* (Anthrax Toxin Receptor 1) and RASGRF2 (Ras Protein Specific Guanine Nucleotide Releasing Factor 2). When comparing these populations with those from HHT samples, we detected striking differences. The single Mature Vein cluster present in healthy tissues segregated into two distinct clusters in HHT samples. One cluster retained the typical Mature Vein signature and did not show major pathway changes relative to healthy tissues. In contrast, the second cluster exhibited a mixed transcriptional identity, expressing venous markers (*ACKR1* and *NR2F2*) along with non-venous genes (*RGCC* and *NOTCH1*). This mixed profile indicated a loss of venous identity. Moreover, this cluster shows features of endothelial activation, including upregulation of VEGFA (Vascular Endothelial Growth Factor A) and ICAM1 (Intercellular Adhesion Molecule 1). Because this specific population appeared exclusively in the HHT samples, we named it “HHT Vein”. Additionally, we identified three other EC sub-populations that were specifically enriched in the HHT samples: d) Capillaries, e) proliferating ECs, and f) lymphatic ECs. These findings align with studies in *Alk1*-deficient mouse AVMs ^33^, and with the identification of an EC cluster displaying “arterial-lymphatic” properties (Lymphatic vessels) in *Alk1*-null mice ^34^.

Pathway enriched analyses compared to Mature Vein ECs revealed increased activity of metabolic pathways, cell adhesion, leukocyte adhesion, and cellular extravasation in the HHT Vein cluster; increased mitochondrial activity in Capillaries; enhanced cell migration, adhesion, and metabolic activity in Artery; activation of pathways related to cell-cycle in the Proliferating Vessel cluster, and up-regulated ECM (extracellular matrix), adhesion and cytoskeleton-related pathways in Lymphatic Vessel cluster (Figure 2E). To better understand pathways activation across EC cluster, we analyzed gene regulation driven by specific transcriptional factors (TFs) by performing regulon analysis, further highlighting the distinct features of the HHT Vein population (Figure 2F). Interestingly, we identified a unique combination of active and inactive transcriptional regulators in the HHT Vein compared to the Mature Vein, including *NR4A2* (Nuclear Receptor Subfamily 4 Group A Member 2), *FOXF1* (Forkhead Box F1), *TEF* (Thyrotroph Embryonic Factor), *GATA6* (GATA Binding Protein 6) and EGR2 (Early Growth Response 2) as activated regulons, suggesting a hybrid and dysregulated endothelial state. Importantly, *FOXF1* and *GATA6* are often re-activated during pathological conditions ^35,36^.

Taken together, these results support findings from multiple AVM mouse models ^33,34,37,38,24,39^ and human brain AVM studies ^40,41^. Furthermore, our analysis demonstrates that telangiectasia contain multiple endothelial subtypes, including ECs with normal venous identity (Mature Vein) and ECs with pathological features (HHT Vein). This distinction strongly suggests that not all ECs within HHT lesions are pathological; rather, a subset undergoes disease-driven reprogramming associated with endothelial activation, loss of identity, and dysfunctional signaling.

### Gene expression characterization of a specific HHT endothelial cluster

The identification of a distinct HHT Vein population, resembling EC clusters described in other vascular malformations ^42^, prompts us to define the molecular differences between the two endothelial states present in HHT telangiectasias: the Mature Vein population, which retains a clear venous identity, and the HHT Vein population, which appears to have partially lost this identity. The HHT Vein cluster expressed genes associated with endothelial activation and features reported in other vascular malformations. These included the pro-angiogenic marker *VEGFA* ^43,44^, the cell cycle inhibitor *CDKN1B* (Cyclin Dependent Kinase Inhibitor 1B) ^45,46^, and inflammatory markers *ICAM1* and *SELE* (Selectin E) ^47–50^. Notably, *PIEZO1* (Piezo-Type Mechanosensitive Ion Channel Component 1) and *KLF4* (Krüppel-like factor 4), two markers previously associated with pathogenic EC states in an HHT mouse model ^24^, were also strongly expressed in this cluster (Figure 2D). To further characterize this aberrant endothelial population, we compared the top up- and down-deregulated genes (DEGs) between Mature Vein and HHT Vein ECs (Figure 3A) and analyzed the pathways associated with these DEGs (Figure 3B). The HHT Vein cluster showed enrichment of pathways involved in response to hormone stimuli, Wnt and TGFβ signaling, cell-substrate adhesion, endothelial development and differentiation, and actin cytoskeleton regulation (Figure 3B). Together, these findings indicate that HHT Vein ECs undergo extensive transcriptional reprogramming and activate pathways known to promote vascular instability.

**Figure 3.**
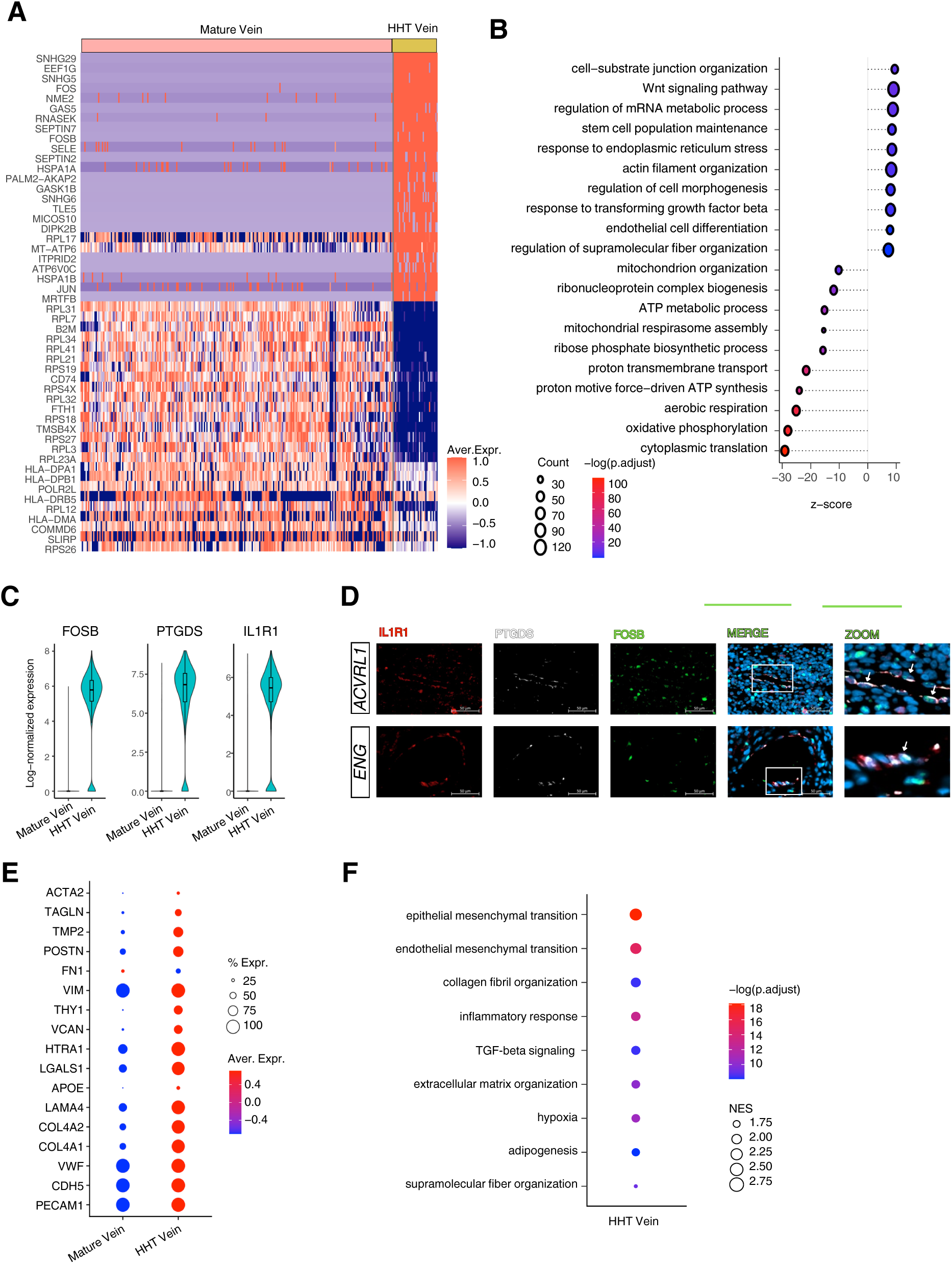
Endothelial cells from HHT Vein sub-populations. **A**, Heat map displaying the most differentially expressed genes (DEGs) in Mature Vein and HHT Vein. Color scale: red, high expression; blue, low expression. **B**, Lollipop plot showing the most active pathways in HHT Vein compared to Mature Vein. Color scale: red, high expression; blue, low expression. **C**, Violin plots comparing the expression levels of FOSB, PTGDS and IL1R1 between Mature Vein and HHT Vein. **D**, Representative images from multiplexing immunofluorescence of FOSB, PTGDS and IL1R1 in *ACVRL1* and *ENG* biopsies. Red, IL1R1; White, PTGDS; Green, FOSB. DAPI, for nuclei. White squares represent the zoomed images. **E**, Dot plot showing expression levels from common genes associated with endothelial-to-mesenchymal transition (EndoMT) in Healthy and HHT Vein cluster. **F**, Dot plot showing pathways associated to EndoMT in Healthy and HHT Vein cluster.

To validate the presence of this pathological EC state *in vivo*, we selected three genes from the top HHT Vein-enriched DEGs with strong relevance to HHT biology: *FOSB* (FosB Proto-Oncogene, AP-1 Transcription Factor Subunit), implicated in angiogenesis ^51,52^; *PTGDS* (Prostaglandin D2 Synthase), associated with inflammation and vasodilation ^53–55^; and *IL1R1* (Interleukin 1 Receptor Type 1), a known amplifier inflammatory signaling ^56^ (Figure 3C and S1B). Using multiplex immunostaining, we confirmed the co-expression of FOSB, PTGDS and IL1R1 in a subset of ECs within telangiectasias (Figure 3D). This co-expression pattern supports the existence of an inflamed, activated, and vasoactive endothelial state, consistent with dysregulated angiogenesis and increased vessel fragility.

Because EC activation, inflammation and remodeling are known to promote the recruitment and activation of myofibroblasts, which contribute to ECM accumulation and fibrosis in damaged tissues ^57–59^, we next asked whether the HHT Vein population also displayed features of EndoMT. Compared with the Mature Vein cluster, HHT Vein ECs showed increased expression of endothelial markers (*PECAM1*, *CDH5* and *VWF*) together with numerous genes associated with EndoMT ^60^, including *ACTA2*, *TAGLN*, *TPM2*, *POSTN*, *VIM*, *THY1*, *VCAN*, *HTRA1*, *LGALS1*, *APOE*, *LAMA4*, *COL4A2*, and *COL4A1* (Figure 3E). Gene Ontology (GO) analyses further revealed enrichment of pathways related to collagen fibril organization (GO:0030199), ECM organization (GO:0030198) and supramolecular fiber organization (GO:0097435) (Figure 3F).

Taken together, these findings demonstrate that dysfunction of the TGFβ/BMP signaling pathways, primarily caused by genetic defects in *ACVRL1* and *ENG*, together with additional second-hit events, drives ECs toward a mesenchymal-like, myofibroblast-associated phenotype. This pathological endothelial state likely contributes to chronic inflammation, fibrotic, and vascular instability in HHT telangiectasias.

### Vessel distribution and fibrotic status of HHT telangiectasias

To better contextualize the molecular alterations identified in HHT ECs, we next examined the tissue-level architecture of telangiectasias and assessed the structural abnormalities associated with AVMs. Nasal telangiectasias from 28 *ACVRL1*-mutant and 5 *ENG*-mutant patients were analyzed using a combination of immunohistochemistry (IHC) and immunofluorescence (IF), followed by computer-assisted image analysis. Hematoxylin and eosin (H&E) staining revealed large, aberrant vascular structures characteristic of AVMs ^61–65^ (Figure S2A). To assess vessel morphology, we performed co-staining with CD34 and ACTA2 to visualize ECs and SMCs, respectively, enabling quantification of vessel size, density and distribution. We first identified the CD34^+^ ACTA2^+^ vessels and classified them into small and large categories (Figure 4A). Both, *ACVRL1*- and *ENG*-mutant samples displayed a higher number of small vessels compared with large ones (Figure 4B and C, Figure S2C and D). However, *ACVRL1* biopsies contained a significantly greater overall number of vessels than *ENG* samples. To determine whether vascular spatial organization deviated from complete spatial randomness, we calculated the NNI for both vessels populations. No significant differences in spatial vascular organization were observed between *ACVRL1* to *ENG* samples (Figure 4D and E). In contrast, within each genotype, small vessels exhibited a clustered distribution (NNI approaching 0), whereas large vessels were more dispersed (NNI ∼1.7) (Figure 4F–G).

**Figure 4.**
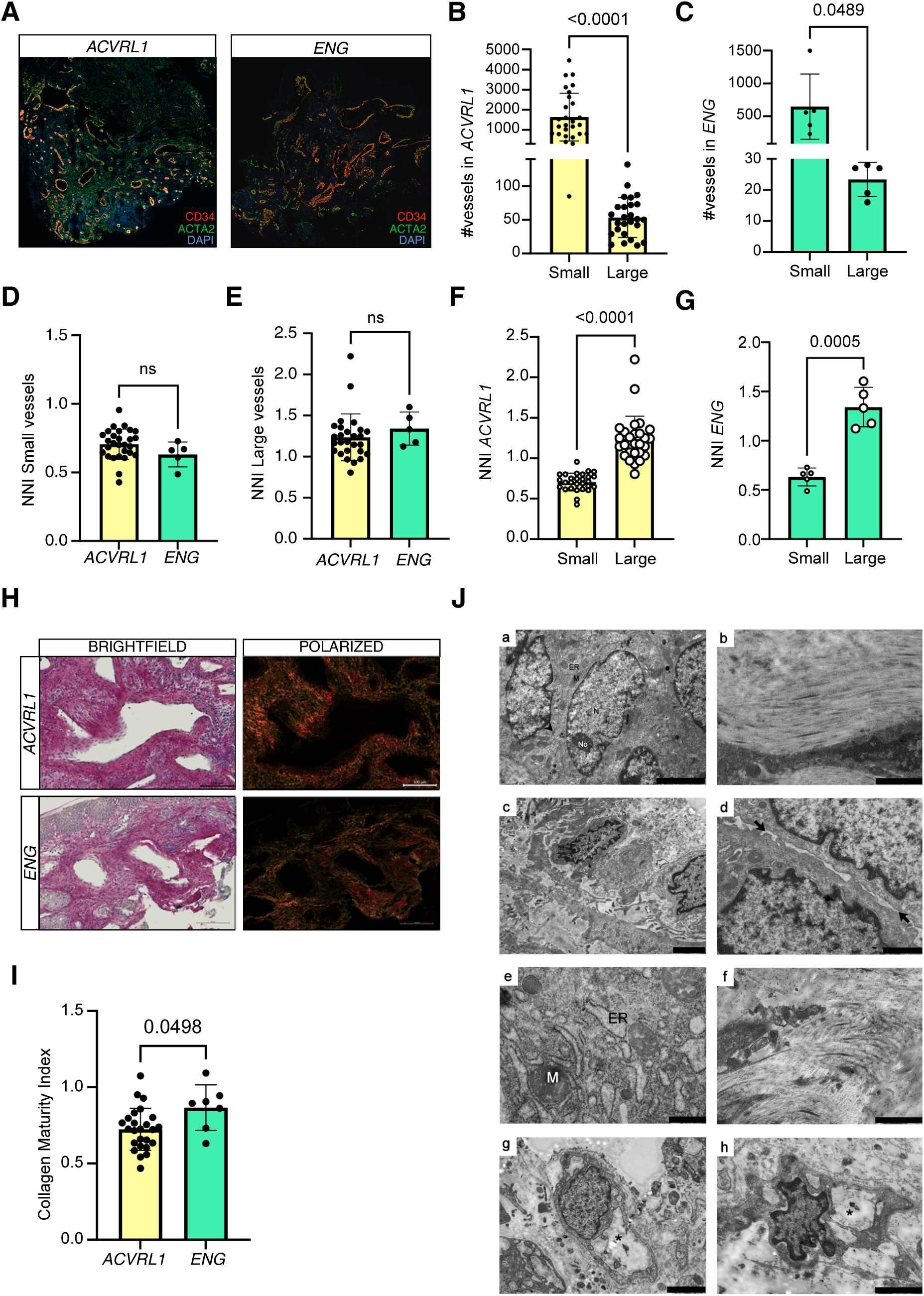
Architectural structures and composition of HHT telangiectasias. **A**, Representative images from *ACVRL1* (left) and *ENG* (right) samples co-stained with CD34 to identify ECs and ACTA2 for SMCs. In blue, DAPI signal identifying nuclei. **B**, Quantification of the number of small and large vessels (CD34^+^ACTA2^+^) counted within *ACVRL1* biopsies. **C**, Quantification of the number of small and large vessels (CD34^+^ACTA2^+^) counted within *ENG* biopsies. **D** and **E**, Quantification of the Nearest Neighbor Index (NNI) comparing: **D**, small vessels within *ACVRL1* and *ENG*; **E**, large vessels within *ACVRL1* and *ENG*; **F**, small and large vessels in *ACVRL1*; and **G**, small and large vessels in *ENG*. **H**, Representative images from Sirius Red staining to visualize collagen deposition in *ACVRL1* (top) and *ENG* (bottom). Tissue structure is visualized under brightfield illumination, whereas polarized light imaging depicts collagen matrix organization and remodeling, with collagen type I appearing as yellow or red fibers and collagen type III as green fibers. **I**, Quantification of the collagen maturity index between *ACVRL1* and *ENG*. **J**, Ultrastructure features by transmission electron microscopy from *ACVRL1* biopsies: (a) Epithelial region showing preserved cells and cellular organelles (M: mitochondria, ER: endoplasmic reticulum, N: nucleus, No: nucleolus), as well as intercellular junctions (arrowhead: desmosomes) ensuring close cell-cell contact. Scale bar: 2µm. (b) Collagen fibrils showing ordered alignment and spatial orientation. Scale bar: 1µm. (c) HTT lesion exhibiting enlarged extracellular space typical of edematous tissues. Scale bar: 2µm. (d) High-magnification micrograph highlighting loss of plasma membrane contact (arrows) in lesioned regions. Scale bar: 1µm. (e) Nanoscale visualization of subcellular stress alterations in pericytes: mitochondria exhibited abnormal matrix density and cristae organization in terms of distribution and abundance, while the rough endoplasmic reticulum appeared enlarged. Scale bar 500nm. (f) Collagen fibrils with diverse spatial orientation accumulated within the extracellular matrix. Cellular debris is visible in close vicinity. Scale bar: 1µm. (g-h) Various degrees of capillary ultrastructural integrity, showing altered endothelial cell preservation and nuclear morphology (asterisk: lumen). Scale bars: 2µm (g), 1µm (h).

Given the transcriptional enrichment of EndoMT and EC-remodeling pathways in HHT Vein ECs, we next investigated fibrosis and collagen deposition in these biopsies. Masson’s Trichrome staining revealed marked collagen accumulation in multiple regions (Figure S2B). Sirius Red staining further distinguished mature collagen type I from newly synthesized collagen type III, revealing area of well-organized collagen alongside regions of dense, disorganized deposition (Figure 4H). Quantification showed a slightly increase in the collagen maturity index in *ENG*-biopsies compared with *ACVRL1* samples (Figure 4I). These findings are consistent with previous reports of excessive collagen deposition contributing to tissue stiffness and pathological remodeling in vascular lesions ^16^.

To further explore these structural alterations at higher resolution, we performed transmission electron microscopy (TEM) (Figure 4J and Figure S2E). Analysis of one *ACVRL*1 and one *ENG* biopsy revealed ultrastructural heterogeneity, with preserved regions interspersed with areas of severe alteration. Intact regions displayed normal epithelial organization with preserved intercellular junctions and continue membranes between adjacent cells (Figure 4J, panel a; Figure S2E, panel i), as well as densely packed, well-aligned collagen fibrils indicative of preserved ECM architecture (Figure 4J, panel b). In contrast, altered regions exhibited features consistent with tissue damage, including disruption of cytoplasmic membranes, particularly in edematous zones, suggesting compromised membrane protein integrity (Figure 4J, panel c, d; Figure S2B, panel j, k). Additional signs of cellular stress were observed, including abnormal mitochondrial and endoplasmic reticulum morphology, notably in pericytes (Figure 4J, panel e). These regions also showed prominent accumulation of collagen fibrils within the ECM (Figure 4J, panel f; Figure S2E, l, m). Similar heterogeneity was observed at the vascular level, where ECs within capillaries displayed both preserved and abnormal ultrastructure features (Figure 4J, panel g, h; Figure S2E, panel n), supporting a spatial association between vascular alterations and tissue damage.

Taken together, these histological and ultrastructural analyses reveal a spatially heterogeneous and structurally compromised microenvironment in HHT telangiectasias. The coexistence of preserved and severely altered regions mirrors the transcriptional heterogeneity observed in EC populations and highlights the strong contribution of fibrosis, ECM remodeling, and endothelial dysfunction to HHT pathology.

### Midkine is involved in endothelial interactions in human HHT nasal telangiectasias

EC communication with surrounding cell types is essential for vascular stability and tissue homeostasis^66,67^. Having identified a pathological EC cluster (HHT Vein ECs), we next sought to determine how these altered ECs interact with other cell populations. To this end, we compared ligand expression between Mature Vein and HHT Vein ECs to identify dysregulated cellular communication elements associated with defective HHT endothelium. We performed cell-cell communication analysis using the Cellchat algorithm to reconstruct global interaction networks in healthy and HHT samples by linking ligands (sender cells) to their cognate receptors (receivers) expressed in other populations (Figure 5A). We then focused specifically on interactions among ECs and non-immune cells (CD45^-^) populations, including fibroblasts and mural cells. Mature Vein ECs from healthy donors exhibited prominent signaling through ligands such as *MDK* (Midkine), *GAS6* (Gamma-carboxyglutamic acid (Gla)-containing protein) and *ESAM* (Endothelial Cell Adhesion Molecule). In contrast, HHT Vein ECs displayed a marked increase in both the number and diversity of interactions (Figure 5B). These cells preferentially signaled through basement membrane components (*COL4A1, COL4A2, LAMA4, LAMB1, HSPG2, HBEGF*), adhesion molecules (*SELE, MPZL1*) and cell-cell communication mediators (*FLRT2, THBS1*), consistent with active ECM remodeling or assembly. They also showed increased expression of stress-response and protective molecules (*CD46, CD55, NAMPT, PPIA* and *EDN1)* and NOTCH ligands (*JAG1*, *JAG2*, *DLL4*). Notably, several ligands highly expressed in Mature Vein ECs were down-regulated in the HHT Vein ECs, indicating loss of key factors required for vascular stability. Among these, MDK emerged as a particularly relevant. MDK is a pro-angiogenic cytokines ^68^ implicated in inflammatory diseases ^69,70^ and vascular homeostasis, including communication with SMCs ^71^. MDK has also been associated with cardio- and neuro-protective effects ^72–74^ and has been observed in brain AVMs ^40,75^. To investigate the role of MDK in this context, we first silenced *ACVRL1* and *ENG* in human umbilical vein EC (HUVEC) to model HHT *in vitro* (Figure S3A and B) and observed a significant reduction in MDK expression (Figure 5C). We then directly silenced MDK (siMDK) in HUVEC (Figure S3C and D), and assessed EC function. MDK knockdown resulted in reduced EC proliferation (Figure S3E), migration (Figure S3F) and angiogenesis capacities (Figure S3G), as assessed respectively by proliferation curve, scratch assay and matrigel tube-forming assay, compared with control cells transduced with scrambled oligos (siSCR). To further characterize the molecular consequences of MDK loss, we performed bulk RNA-seq from HUVEC on siSCR and siMDK HUVECs. Although numerous genes were differentially expressed (Figure S3H and File S2), no single pathway reached strong statistical enrichment, consistent with the pleiotropic role of MDK ^69–71^. Down-regulated genes in siMDK cells-including *LSS*, *ACAT2*, *PGAM2*, *AKT1*, *PHKG1*, *DVL3*, *ATP6V1C2*, *PLAT*, *MASP2* and *BCAN*- are associated with lipid metabolism, cholesterol biosynthesis, VEGF signaling, hypoxia and angiogenesis. Conversely, up-regulated genes such as *PTGS2*, *ASCC3*, *VCAN*, *ITGAV*, *BMPR2*, *SMAD5*, *TEAD1*, *BIRC2*, *SEMA3C* are linked to BMP/TGFβ signaling, PlexinD1 signaling, eicosanoids pathways, and Carbohydrate Sulfotransferases. Moreover, the presence of several non-annotated genes may indicate that MDK-related transcriptional changes involve poorly characterized genomic regions, possibly including epigenetic regulatory mechanisms, consistent with previous reports ^76^.

**Figure 5.**
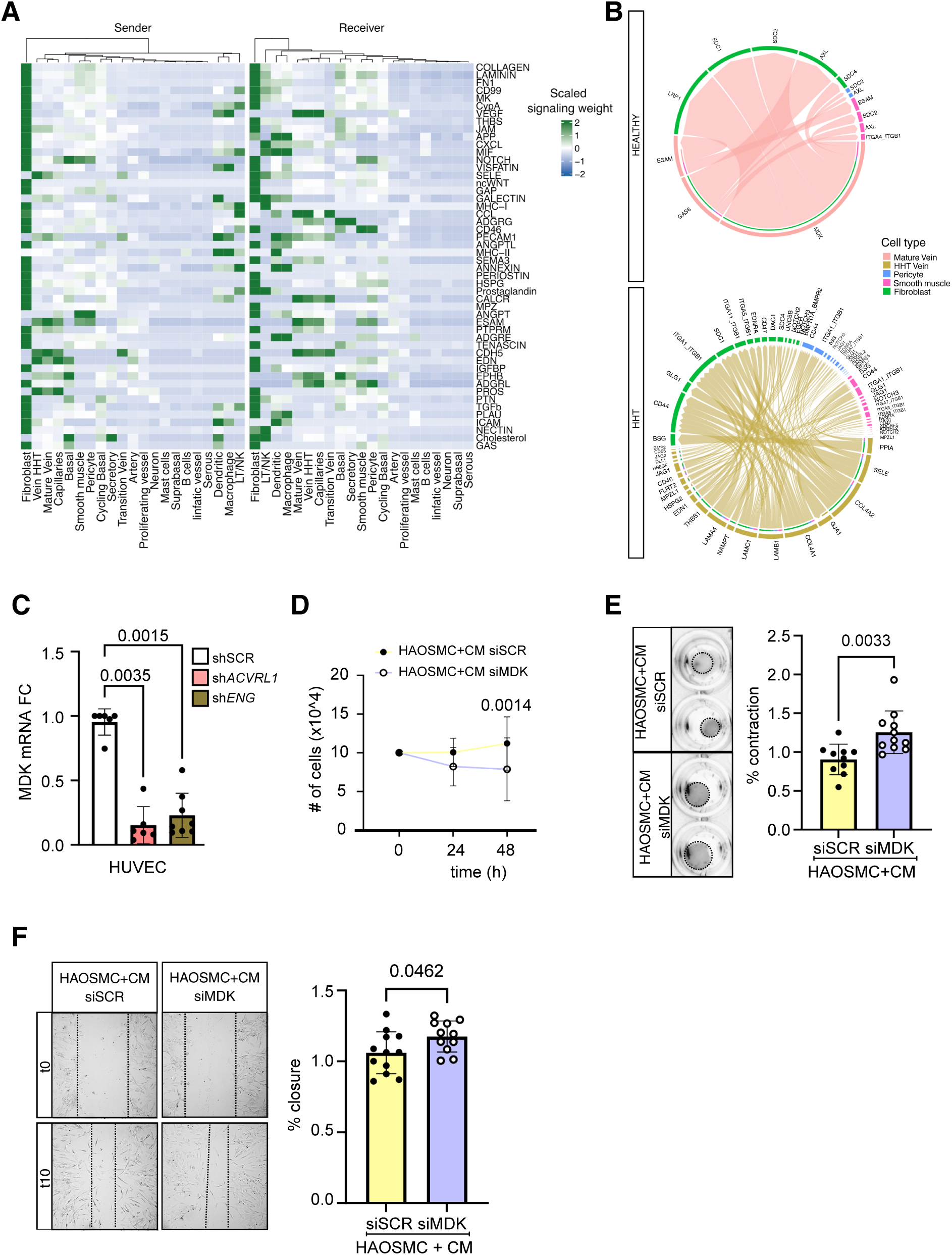
Cellular interactions between ECs and non-immune cells in HHT and healthy samples. **A**, Heat map of the activated signaling pathways on ECs pinpointing the contribution of the different cell types (CD45^-^). Sender: produced ligands by the different cell types that activate ECs. Receiver: receptors expressed in the different cell types and that activate ECs. **B**, CellChat showing the interactions between ligands produced by ECs and active receptors in CD45^-^ cells, including fibroblasts, smooth muscle cells and pericytes in Healthy samples (upper) compared to HHT samples (lower). **C**, Midkine (MDK) mRNA Fold change (FC) from qRT-PCR performed in HUVEC silenced for *ACVRL1* and *ENG*. Phenotypic assays performed in HAOSMC with conditioned medium (CM) from HUVEC siSCR and siMDK: **D**, Proliferation curve from Cells plated (t0) and counted after 24 and 48 hours. **E**, Representative pictures from contraction assay and corresponding histogram of the quantification. **F**, Representative pictures from migration assay and corresponding histogram of the quantification.

Given the close functional relationship between ECs and SMCs in this pathological context, as support by the CellChat interaction analysis, and the observed loss of MDK-mediated signaling in HHT Vein ECs, we next assessed the impact of EC-derived MDK on SMC behavior. SMCs exposed to conditioned medium (CM) from EC-siMDK (Figure S3C) showed reduced proliferation (Figure 5D) and contractile capacity (Figure 5E), along with a slightly no statistically reduction in the expression of differentiation markers (Figure S3I). Migratory capacity was observed to be increased in SMCs exposed to CM of EC-siMDK (Figure 5F).

Collectively, these findings demonstrate that extensive molecular reprogramming occurring in HHT Vein ECs leads to profound alterations in cell–cell communication. In particular, loss of MDK signaling impairs EC function and disrupts EC–SMC interactions, contributing to abnormal ECM dynamics, defective vessel stabilization, and pathological microenvironment characteristic of HHT telangiectasias.

### Spatial transcriptomics localized the HHT specific endothelial cluster in human HHT nasal telangiectasias

To validate the presence of HHT Vein population within the patients’ telangiectasia and to uncover cellular crosstalk, we performed Spatial Transcriptomics on one *ACVRL1*-biopsy and one *ENG*-biopsy. From each section, after quality filtering, we captured 38051 and 15684 cells, respectively. Unsupervised clustering identified 14 distinct populations (Figure S4A-B), which were annotated using known gene signatures. In both biopsies, we identified clusters enriched in vascular gene signatures and used those cells to dissect the vascular heterogeneity (Figure S4C). Cells of interest were subjected to a re-clustering analysis; veins, arteries and capillaries sub-clusters were identified using our scRNA-seq as well as public gene signatures ^77^. From the re-clustering of vascular cells, we obtained 14 populations with distinct EC signatures in *ACVRL1* (Figure 6A-B) and *ENG* (Figure 6C-D). We focused on populations reflecting venous identity and recognized Cluster 2 in *ACVRL1*-biopsy (Figure 6B) and Cluster 6 in *ENG*-biopsy (Figure 6D) as candidate clusters corresponding to the HHT Vein population identified by scRNA-seq. To better resolve EC sub-populations, we further re-clustered ECs within Cluster 2 from the *ACVRL1*-biopsy at a resolution of 0.3, identifying five sub-clusters (Figure 6E-F), and ECs within Cluster 6 from the *ENG*-biopsy at a resolution of 0.6, identifying six EC sub-populations (Figure 6G-H). Integration with molecular signatures from our scRNA-seq dataset enabled precise annotation of EC subsets in both *ACRVL1*- (Figure S4D) and *ENG*-biopsies (Figure S4E). In the *ACVRL1*-biopsy, sub-clusters 0 and 3 displayed venous identity based on the expression of canonical markers, such as *ACKR1*. Sub-cluster 0 showed a “quiescent” profile, resembling the Mature Vein population, whereas sub-cluster 3 exhibited an “activated” phenotype characterized by elevated expression of *VCAM1*, *ICAM1* and *SELE*, consistent with endothelial dysfunction. Accordingly, sub-cluster 3 was annotated as the HHT Vein equivalent.

**Figure 6.**
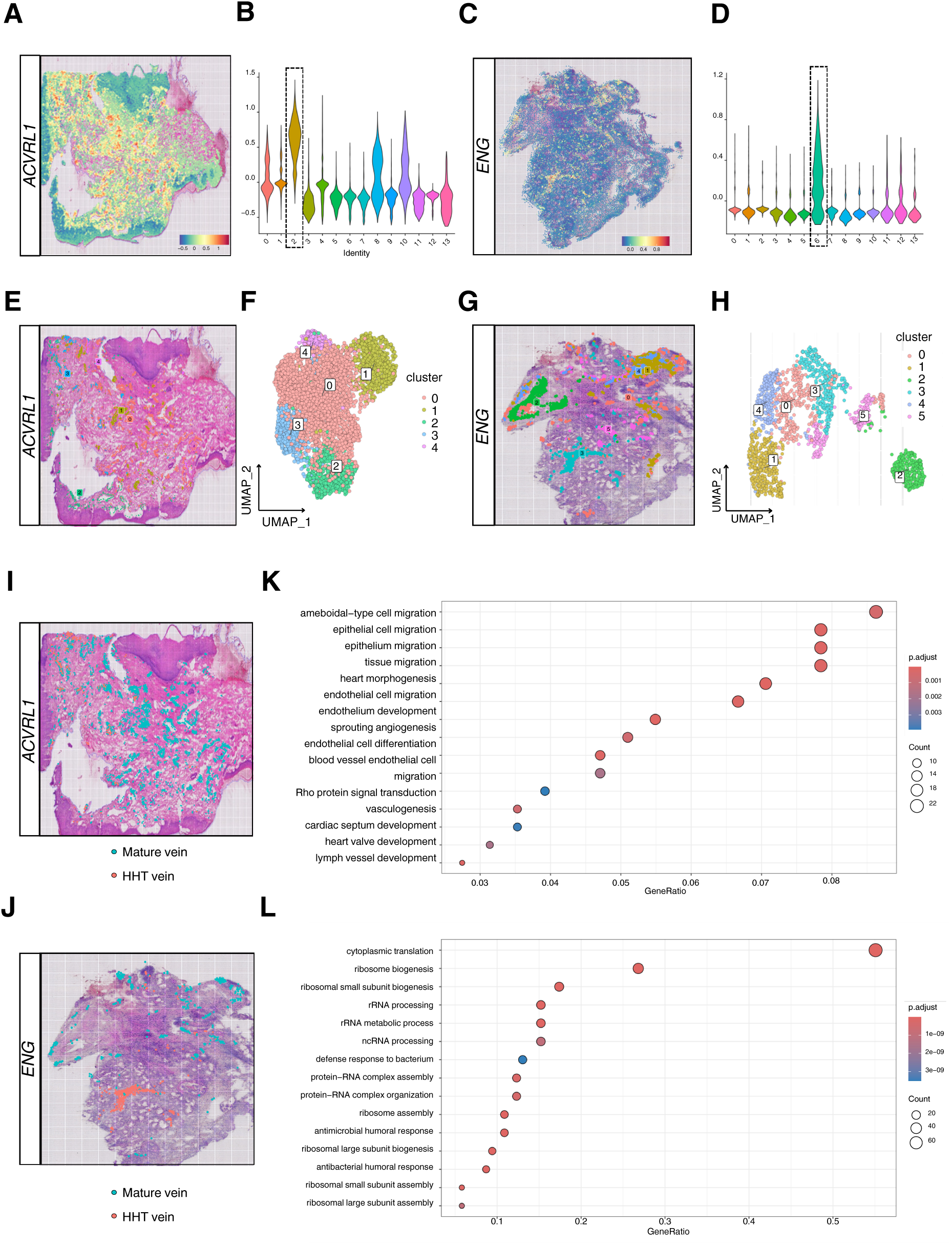
Identifying of the HHT Vein population through Spatial Transcriptomics. EC cluster identification through re-clustering of all vascular components. **A** and **B**, DIM plot and violin plot after re-clustering in *ACVRL1*-biopsy. **C** and **D**, DIM plot and violin plot after re-clustering in *ENG*-biopsy. Dashed lines identify the endothelial cells cluster. Re-clustering performed to identify the different EC subsets. **E** and **F**, DIM plot and UMAP showing EC populations from Cluster 2 at a 0.3 of resolution in *ACVRL1*-biopsy. **G** and **H**, DIM plot and UMAP showing EC populations from Cluster 6 at a 0.6 of resolution in *ENG*-biopsy. **I** and **J**, Spatially visualization of the Mature and HHT Vein clusters in *ACVRL1*-biopsy and *ENG*-biopsy. **K** and **L**, GO annotations showing up-regulated pathways in HHT Vein EC compared to Mature Vein EC in *ACVRL1*-biopsy and *ENG*-biopsy. Color scale: red, high expression; blue, low expression.

Sub-cluster 1 exhibited an arterial signature (*SEMA3G*, *GJA5*, *NOTCH1*, *HEY1/2*); sub-cluster 2 represented capillaries (high expression of *RGCC*); and sub-cluster 4 appeared to represent a transition population, expressing smooth muscle-associated genes (*ACTA2*, *KCNJ8*, *MYH11*), potentially reflecting EndoMT-related states. Similarly, in the *ENG*-biopsy, sub-clusters 0, 3 and 4 were also recognized as venous ECs, with sub-clusters 0 and 4 showing a “quiescent” profile (Mature Vein) and sub-cluster 3 displaying activated EC features (HHT Vein). Sub-clusters 1 and 2 exhibited a SMC-like gene profile, while sub-cluster 5 showed artery identity. After defining EC subtypes and identifying and identified the populations of interest, we visualized the spatial localization of Mature and HHT Vein cells across the tissue sections in both *ACVRL1* (Figure 6I) and *ENG* (Figure 6J) biopsies, and evaluated their transcriptomic differences. Compared with Mature Vein cells, the HHT Vein population showed up-regulated genes related to vascular development in the *ACRVL1*-biopsy (Figure 6K) and genes involved in RNA processing and biogenesis in the *ENG*-biopsy exhibited (Figure 6L).

Overall, these results validate the presence of the HHT Vein population spatially within telangiectasias and suggest the coexistence of different EC populations with venous characteristics that diverge from a stable physiological state towards a more dynamic and less defined identity.

### Neighborhood analysis revealed active remodeling surrounding HHT ECs in telangiectasias

Given the extensive dysregulation of EC communication observed in the scRNA-seq dataset, we hypothesized that cells physically in contact with Mature Vein ECs would differ from those neighboring HHT Vein ECs. Thus, we performed spatial identification of all neighboring cells in *ACVRL1* biopsy around Mature Vein (Figure 7A-B) and HHT Vein ECs (Figure 7C-D), as well as in the *ENG* biopsy (Figure S4F-I). Consistent trends were observed in the *ENG* biopsy, although reduced cellular recovery limited the depth of downstream analyses. Overall, without distinguishing between Mature or HHT EC subtypes, keratinocytes were frequently detected around vessels, indicating a local inflammatory and wound-healing environment ^78,79^. We then specifically compared the cellular microenvironment of Mature Vein *vs* HHT Vein ECs at the transcriptomic level. Across both contexts, fibroblasts emerged as the predominant interacting cells, expressing ECM- and fibroblast-associated markers, such as *COL1A1*, *COL3A1*, *FBLN1* and *MMPs*. Strikingly, fibroblasts surrounding HHT Vein ECs showed activation of pathways linked to myofibroblasts differentiation, ECM remodeling, angiogenesis, inflammation and metabolic stress, while fibroblasts near to Mature Vein ECs maintained pathways related to epithelial differentiation and proliferation (Figure 7E).

**Figure 7.**
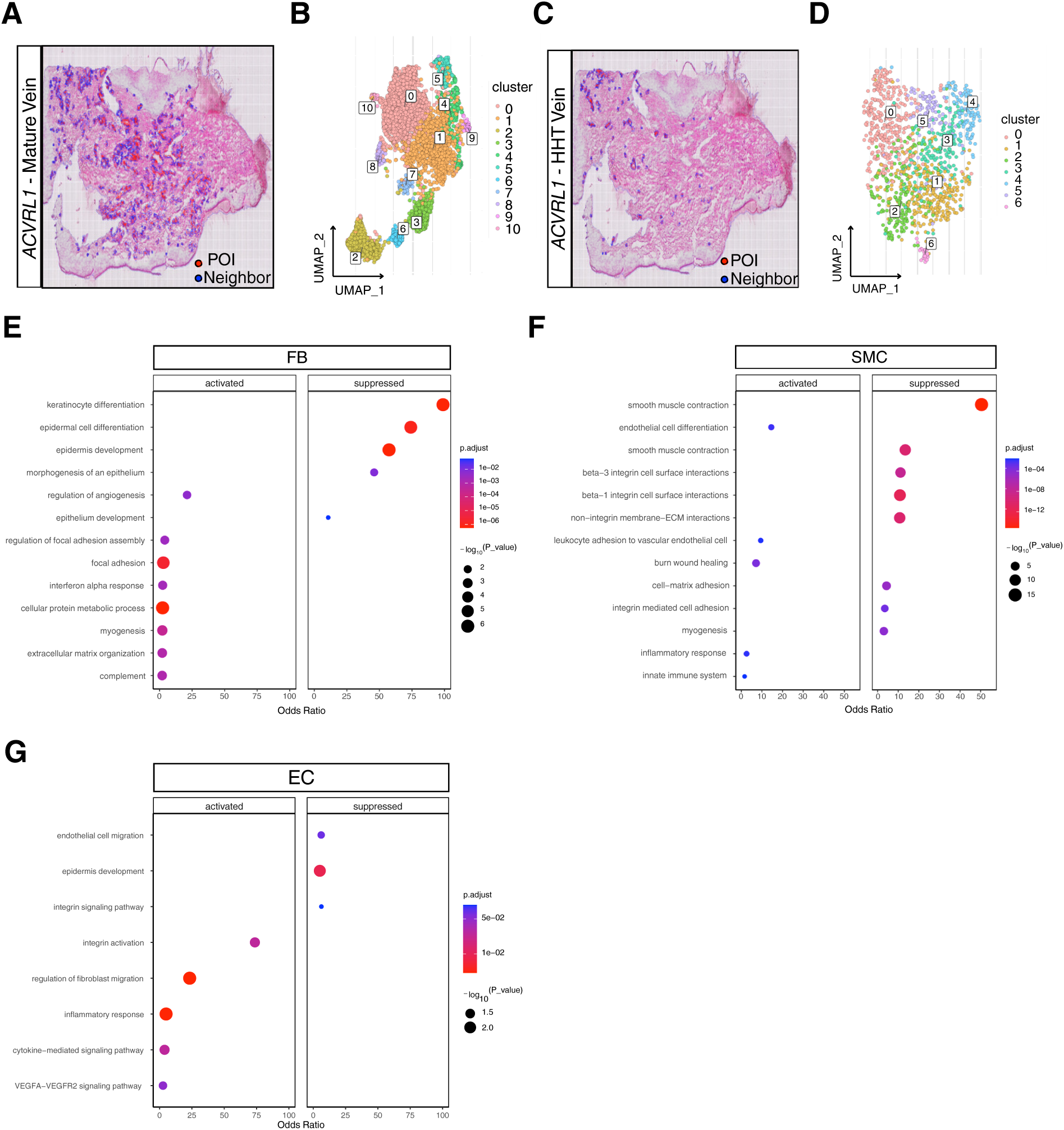
Cellular neighbors surrounding Mature and HHT Vein clusters. Cells recognized in *ACVRL1*-biopsy within 100μm of distance from selected ECs (POI: point of interest) and identified as neighbors: **A** and **B**, DIM plot and UMAP plot identifying neighbors from Mature Vein EC; **C** and **D**, DIM plot and UMAP plot revealing neighbors from HHT Vein ECs. Color code: POI, identifying ECs (red) and identified neighbor cells (blue). UMAP represent the re-clustering of all neighbor cells identified. Activated and Suppressed pathways in neighbor cells from HHT Vein compared to Mature Vein from *ACVRL1* biopsy: **E**, Fibroblasts (FB); **F**, Smooth muscle cells (SMC); **G**, endothelial cells (ECs).

As expected, SMCs (*ACTA2, MYH11, ACTG2*) were also detected in the proximity of ECs. Interestingly, SMCs adjacent to HHT Vein ECs exhibited a profile characterized by increased ECM remodeling, oxidative stress and inflammatory pathways (Figure 7F). In contrast, contractile and adhesion genes were down-regulated, indicating loss of vascular support and stability. Additionally, endothelial populations were found adjacent to the vessels. These is likely due to tissue heterogeneity. While Mature and HHT Vein ECs are the dominant populations, the adjacent ECs likely represent functional states influenced by the local microenvironment. Indeed, surrounding HHT Vein, we observed ECs with an activated profile, whereas ECs surrounding Mature Vein displayed a more mature and quiescent phenotype (Figure 7G).

Altogether, these results suggest increased cellular remodeling in proximity to active and more pathogenic ECs, indicating a direct impact of cellular communication between ECs and other vascular cells, ultimately contributing to overall vascular instability.

## DISCUSSION

Vascular development, remodeling and maturation are tightly regulated processes. During embryonic and post-natal development, blood flow drives vascular remolding, enabling the formation of morphologically, transcriptionally and functionally distinct arterial, venous, and capillary networks ^33,80^. HHT is characterized by malformed vascular connections that bypass capillaries, forming direct arteriovenous shunts known as telangiectasias or AVMs ^1^.

In this study, we used nasal telangiectasia biopsies from HHT patients undergoing surgery to dissect the cellular and molecular landscape of human HHT lesions. By integrating scRNA-seq, spatial transcriptomics, TEM, and histological analyses, we provided a direct molecular characterization of diseased HHT tissues, offering an *ex vivo* snapshot of the transcriptional and structural alterations underlying telangiectasia formation.

Recently, significant efforts have led to the identification of distinct EC populations within AVMs in mouse models of HHT ^33,34,37,38,24,39^ as well as human brain AVMs ^41,75^, revealing potential pathological populations driving AVM development. In line with these findings, our scRNA-seq data revealed substantial transcriptional reprogramming among ECs in HHT telangiectasias. We observed a redistribution of EC clusters, with an increase of lymphatic, proliferative and capillaries ECs in HHT biopsies. Most strikingly, the venous endothelial compartment segregated into two distinct clusters in HHT samples: one maintaining a stable venous identity (“Mature Vein ECs”), and a second exhibiting distinct features of endothelial activation (“HHT Vein ECs”). During physiological angiogenesis, venous ECs can proliferate and differentiate into tip, capillary, and arterial ECs, migrating against blood flow to shape new vasculature ^80^. Disruption of endothelial venous identity can therefore have profound pathological consequences. For instance, venous ECs have been shown to aberrantly connect to arterial ECs during AVM formation ^80^. In addition, recent genetic and preclinical studies have identified disruptions in pathways regulating angiogenesis, vascular stability, and blood flow as key contributors to vascular malformations ^81^.

Consistent with this, the HHT Vein cluster in our dataset expressed a hybrid molecular signature combining venous, arterial, and capillary markers, indicating a profound loss of EC identity. This phenotype suggests that the HHT microenvironment, together with loss-of-function mutations in *ENG* or *ACVRL1*, drives transcriptional reprogramming toward pathological endothelial states. Interestingly, the HHT Vein population express elevated levels of *PIEZO1*, a mechanosensitive ion channel recently implicated in AVM development and aberrant endothelial response to flow^33^. This cluster displayed features of endothelial activation, hormone responsiveness, and dynamic remodeling programs. Secreted factors previously reported to be elevated in HHT, including VEGF and TGFβ ^82–84^, were also enriched in this population. We further observed pronounced activation of TGFβ/BMP signaling within the HHT Vein ECs. Among the transcriptional factors, GATA6 captured our attention due to its high activity. GATA6 is context-dependent and is involved in cell proliferation ^85^, TGFβ modulation ^36^, inflammatory response ^86^, and monocyte adherence and migration ^87,88^. GATA6 has been linked to different vascular pathologies: it is down-regulated in systemic sclerosis-associated PAH ^89^, but up-regulated in atherosclerosis and its inhibition ameliorates plaque development ^87^, suggesting microenviromental cues critically shape its function. In our context, GATA6 activation may stem from impaired ENG/ACVRL1-mediated BMP signaling, contributing to the inflammatory and activated state of HHT Vein ECs.

Moreover, we found co-expression of *IL1R1*, *PTGDS* and *FOSB* in HHT Vein ECs, along with up-regulation of genes associated with EC activation, such as *ICAM1*, *VCAM*, *ESEL*. *IL1R1* is associated with inflammation and its inhibition has been shown to alleviate sepsis-associated encephalopathy ^90^. Moreover, *IL1R1* has been also linked to plaque instability during atherosclerosis and to EndoMT ^91^. The TGFβ pathway activates the *AP-1* transcription factor complex (Jun, Fos and ATF proteins) ^92,93^, and both aberrant TGFβ singling and AP-1 activation are known to promote EndoMT ^94^. *PTGDS* and *FOSB* are both *AP-1-*regulated genes ^51,95,96^ involved in vascular inflammation and shear stress response. Notably, shear stress can induce *AP-1*, promoting *PTGDS* expression, and recent studies have shown that *PIEZO1* - upregulated during shear stress conditions - can enhance prostaglandin production ^96,97^. Altogether, the co-expression of these genes suggests a mechanically driven, pro-inflammatory endothelial state contributing to vascular instability.

The HHT Vein cluster also exhibit increased expression of mesenchymal-like genes, consistent with EndoMT, a process known to drive vascular instability in pathological conditions ^98–100^. Since TGFβ is a primary inducer of EndoMT ^8^, its over-activation in HHT Vein ECs likely contributes to this phenotype. EndoMT is closely linked to fibrosis ^101^ and our transcriptomics analyses aligned with histological observations showing extensive collagen deposition in multiple biopsy regions. TEM analyses further confirmed disorganized collagen and ultrastructure abnormalities. Importantly, tissues exhibited marked heterogeneity, as not all regions within telangiectasias displayed abnormal features. Some areas retained well-preserved vascular morphology, consistent with transcriptionally defined “healthy” Mature Vein and arterial EC clusters. These findings support the idea that HHT pathology arises from a subset of aberrantly reprogrammed ECs rather than a global endothelial dysfunction.

Another important aspect is that vascular maturation depends on recruitment of mural cells – PCs or SMCs – to stabilize the vessel wall and regulate blood flow ^26,33,45,102–104^. In HHT, not only ECs but also mural cells and other stromal populations appear compromised. Recent studies have highlighted mural dysfunction in HHT, as well as in other rare vascular diseases ^105^. Our cell-cell interaction analyses revealed a marked increase in EC-driven interactions in HHT samples compared to healthy controls. Notably, HHT Vein ECs showed increased inflammatory and stress-related signaling, together with a specific loss of MDK, a ligand abundantly expressed in Mature Vein ECs. MDK is a pro-angiogenic cytokine involved in tissue remodeling and inflammatory response. Its down-regulation in HHT Vein ECs, confirmed in our *in vitro ACVRL1-* and *ENG-*silenced models, suggests that MDK depletion may contribute to vascular destabilization.

Finally, using high-resolution spatial transcriptomic, we validated the presence of the HHT Vein population *in situ* and mapped its cellular microenvironment. HHT Vein ECs were consistently surrounded by fibroblasts exhibiting myofibroblast-like signatures, including ECM activation, angiogenesis, inflammation, and metabolic stress, consistent with a pro-fibrotic and remodeling-prone niche. Mural cells adjacent to HHT Vein ECs showed reduced expression of contractile markers and increased dedifferentiation, ECM remodeling, oxidative stress and inflammatory pathways, indicating compromised vessel stabilization. Additional EC subpopulations surrounding HHT Vein ECs were enriched for hypoxia, stress, and angiogenic programs while lacking endothelial integrity markers. Together, these findings reveal a profoundly altered and unstable microenvironment surrounding pathological HHT Vein ECs.

This study provides a comprehensive multi-omic and spatially resolved portrait of HHT telangiectasias, revealing that disease pathology is driven by a specific subset of ECs undergoing identity loss, inflammatory activation, and EndoMT. These pathological ECs arise in the context of disrupted TGFβ/BMP signaling and interact with a reactive microenvironment composed of activated fibroblasts and dysfunctional mural cells. The combined effect of these aberrant cellular states likely promotes matrix remodeling, vascular instability, and AVM formation.

Our findings highlight the biological complexity of HHT and emphasize the need for future studies to refine the cellular and molecular mechanisms driving disease onset and progression. By identifying pathological EC states and microenvironmental interactions that contribute to telangiectasia formation, this work lays the foundation for the development of targeted therapeutic strategies aimed at stabilizing the vasculature and mitigating disease severity.

## Acknowledgments

The authors would really like to thank the patients and their families for the support and willingness of participating to the research project. We thank all the Unit of Otolaryngology and the Unit of SS Genetica Clinica from IRCCS Policlinico San Matteo for their continuous support and collaboration. We also want to thank the Genomic Core Facility HuGe from Humanitas Research Hospital, specially Gianluca Basso and Desiree Giuliano, for the support on the scRNA-seq process. The authors also thank Dr. Massimo Boiocchi from the Transmission Electron Microscopy Facility, Centro Grandi Strumenti (CGS), University of Pavia. Fondazione Humanitas per la Ricerca is acknowledged for supporting MAAAH, a co-author of this study.

## Author contributions

A.S., L.L. and M.C. conceptualized and planned experiments. A.S., L.C. and M.C. performed the experiments. L.L. and R.C. analyzed scRNA-seq and Spatial Transcriptomics data. C.C. and M.C. performed TEM acquisition and analyses. M.H. and F.G. contributed to the histological planning and analyses. S. I. visited and enrolled patients and collected biopsies. F.S. and C.O. processed and provided genetic information from HHT patients. F.P., L.E. and M.C. provided funding. A.S., L.L., L.E. and M.C. wrote and edited the final manuscript with input and approval from all authors.

